# Yolk-Sac-Derived Macrophages Progressively Expand in the Mouse Kidney with Age

**DOI:** 10.1101/811927

**Authors:** Shintaro Ide, Yasuhito Yahara, Yoshihiko Kobayashi, Sarah A. Strausser, Anisha Watwe, Jamie R. Pivratsky, Steven D. Crowley, Mari L. Shinohara, Benjamin A. Alman, Tomokazu Souma

## Abstract

Renal macrophages represent a highly heterogeneous and specialized population of myeloid cells with mixed developmental origins from the yolk-sac and hematopoietic stem cells (HSC). They promote both injury and repair by regulating inflammation, angiogenesis, and tissue remodeling. Recent reports highlight differential roles for ontogenically distinct renal macrophage populations in disease. However, little is known about how these populations change over time in normal, uninjured kidneys. Prior reports demonstrated a high proportion of HSC-derived macrophages in the young adult kidney. Unexpectedly, using genetic fate-mapping and parabiosis studies, we find that yolk-sac-derived macrophages progressively expand in number with age and become a major contributor to the renal macrophage population in older mice. This chronological shift in macrophage composition involves cellular proliferation and recruitment from circulating progenitors and may contribute to the distinct immune responses, limited reparative capacity, and increased disease susceptibility of kidneys in the elderly population.

---

Tissue-resident macrophages constitute a highly heterogeneous and specialized population of myeloid cells, reflecting the diversity of their developmental origins and tissue microenvironments [1-10]. In addition to their critical roles in host defense against pathogens, macrophages are central to sterile inflammation, angiogenesis, and tissue remodeling, making them an attractive target for therapeutic intervention. Tissue-resident macrophages originate from at least three distinct progenitors: (*i*) macrophage colony-stimulating factor 1 receptor (CSF1R)-positive yolk-sac macrophages; (*ii*) CX3C chemokine receptor 1 (CX3CR1)-positive yolk-sac macrophages, also known as pre-macrophages; and (*iii*) embryonic and neonatal hematopoietic stem cells (HSC) [1-9]. These populations are maintained *in situ* by self-renewal, largely independent of adult hematopoiesis [1-8, 10-12].

Most tissues have mixed populations of different ontogenically-derived macrophages, and their relative contributions and temporal kinetics are tissue-specific. For example, microglia, Kupffer cells, and Langerhans cells originate from the yolk-sac, with minimal contribution from HSCs [2-6, 11]. However, macrophages in the intestinal wall are rapidly replaced by HSC-derived macrophages after birth [13]. Importantly, investigators now recognize that macrophage ontogeny contributes to their roles in disease processes such as cancer progression; in pancreatic cancer, for example, yolk-sac-derived macrophages are fibrogenic, while HSC-derived macrophages are immunogenic. This raises the possibility that developmental programs influence how macrophages differentially respond to disease insults [14].

Renal macrophages are found in an intricate network surrounding the renal tubular epithelium [15-17] and have mixed origins from both yolk-sac and HSC [3, 6, 8, 18]. They exert unique functions depending on their anatomical locations; monitoring and clearing immune complexes is a function of cortical macrophages while bacterial immunity is the responsibility of medullary macrophages [15, 16]. Renal macrophages critically control renal inflammation and tissue remodeling after injury with robust phenotypic reprogramming [19]. Despite the importance of these roles, little is known about how the proportion and distribution of macrophages of different ontogeny change over time in the normal kidney and whether this influences the increased susceptibility and poorer outcomes of older patients to acute and chronic kidney diseases [20-22]. While most preclinical models of kidney diseases have used young animals, recent papers highlight distinct immune responses in aged mouse kidneys. Aged kidneys exhibit more severe inflammation than young kidneys in response to ischemic and toxic insults, leading to maladaptive repair and organ dysfunction [22, 23]. Furthermore, there is a growing interest in aging as a fundamental determinant of macrophage heterogeneity, such as in the heart and serous cavities [24-26]. However, data on the effects of aging on the renal macrophage populations are lacking. Here, using complementary *in vivo* genetic fate-mapping and parabiotic approaches, we identify a previously unappreciated increase in the proportion of yolk-sac-derived macrophages in the kidney with age.

## Results and Discussion

Two complementary strategies were used to fate-map CSF1R^+^, and CX3CR1^+^ yolk-sac-derived macrophages (Fig. 1A). These partly overlapping populations appear in the yolk-sac around E8.5. Subsequently, from E9.0 until E14.5, they proliferate, migrate, and colonize the embryo through the vascular system [9, 18]. To lineage-label these cells, we exposed *Csf1r-Mer-iCre-Mer*; *Rosa26-tdTomato* and *Cx3Cr1-CreER*; *Rosa26-tdTomato* embryos to 4-hydroxytamoxifen (4-OHT) at E8.5 and E9.5, respectively (Fig. 1A), [1]. This efficiently and irreversibly labels yolk-sac-derived macrophages with the tdTomato reporter. Importantly, the approach does not label fetal monocytes or HSCs, as we recently determined by single-cell RNA sequencing at E14.5 (Y.Y et al., *in revision*).

**Figure 1.**
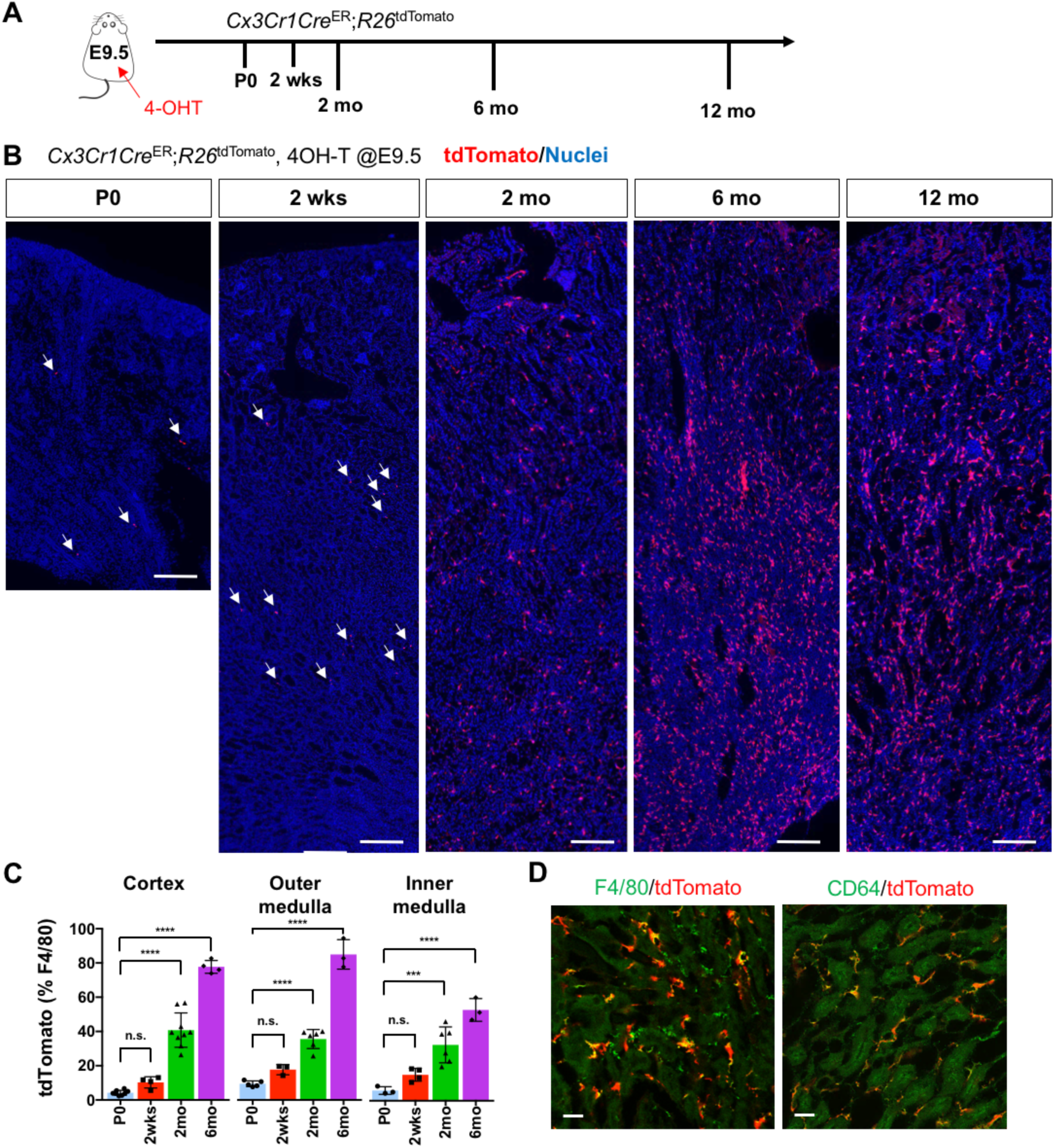
CX3CR1-positive yolk-sac macrophage descendants progressively expand in number in kidneys with age. (A) Fate-mapping strategies of CX3CR1^+^ yolk-sac macrophages. 4-hydroxytamoxifen (4-OHT) was injected once into pregnant dams at 9.5 dpc and offspring analyzed at the indicated times (n=4-6 for P0 to 6-month-old; n=2 for 12-month-old). Yolk-sac macrophages and their progeny are irreversibly tagged with tdTomato. (B) Distribution of CX3CR1-lineage cells in postnatal kidneys. Arrows: CX3CR1-lineage cells. (C) Percentage of tdTomato^+^ to F4/80^+^ cells. Data are represented as means ± S.D. ***, *P*<0.001; ****, *P*<0.0001; n.s., not significant. (D) Confocal images of F4/80 and CD64 staining in aged kidneys (6 mo) with CX3CR1-lineage tracing (n=3). Scale bars: 200 μm in B; 10 μm in D.

At postnatal day 0 (P0), we detected a small number of tdTomato^+^ cells in kidneys from both lines (Fig. 1 and 2). A previous fate-mapping strategy that labels all HSC-derived cells indicated that 40 to 50% of tissue-resident macrophages in the young adult kidney originate from HSC; the remainder was inferred to derive from yolk-sac hematopoiesis [3]. Consistent with this inference, we found CX3CR1-lineage labeled cells in kidneys from birth, with numbers increasing progressively over time (2 weeks, 2 months and 6 months; Fig. 1, B and C). Surprisingly, we observed an unexpected large increase in the proportion of tdTomato-positive cells relative to total F4/80-positive cells at 6 months, especially in the cortex and outer medulla, despite no significant change in the number of F4/80-positive cells per section. This increase was maintained up to 1 year (Fig. 1B). The tdTomato-labeled cells were positive for mature macrophage markers, F4/80 and CD64 (Fig. 1D), [17, 27]. By contrast, we observed only a few F4/80-positive CSF1R-lineage cells inside the kidneys (Fig. 2, B and C), as reported previously [3]. These data demonstrate that CX3CR1^+^ yolk-sac macrophages and their descendants are major contributors to the resident renal macrophage population in aged kidneys.

**Figure 2.**
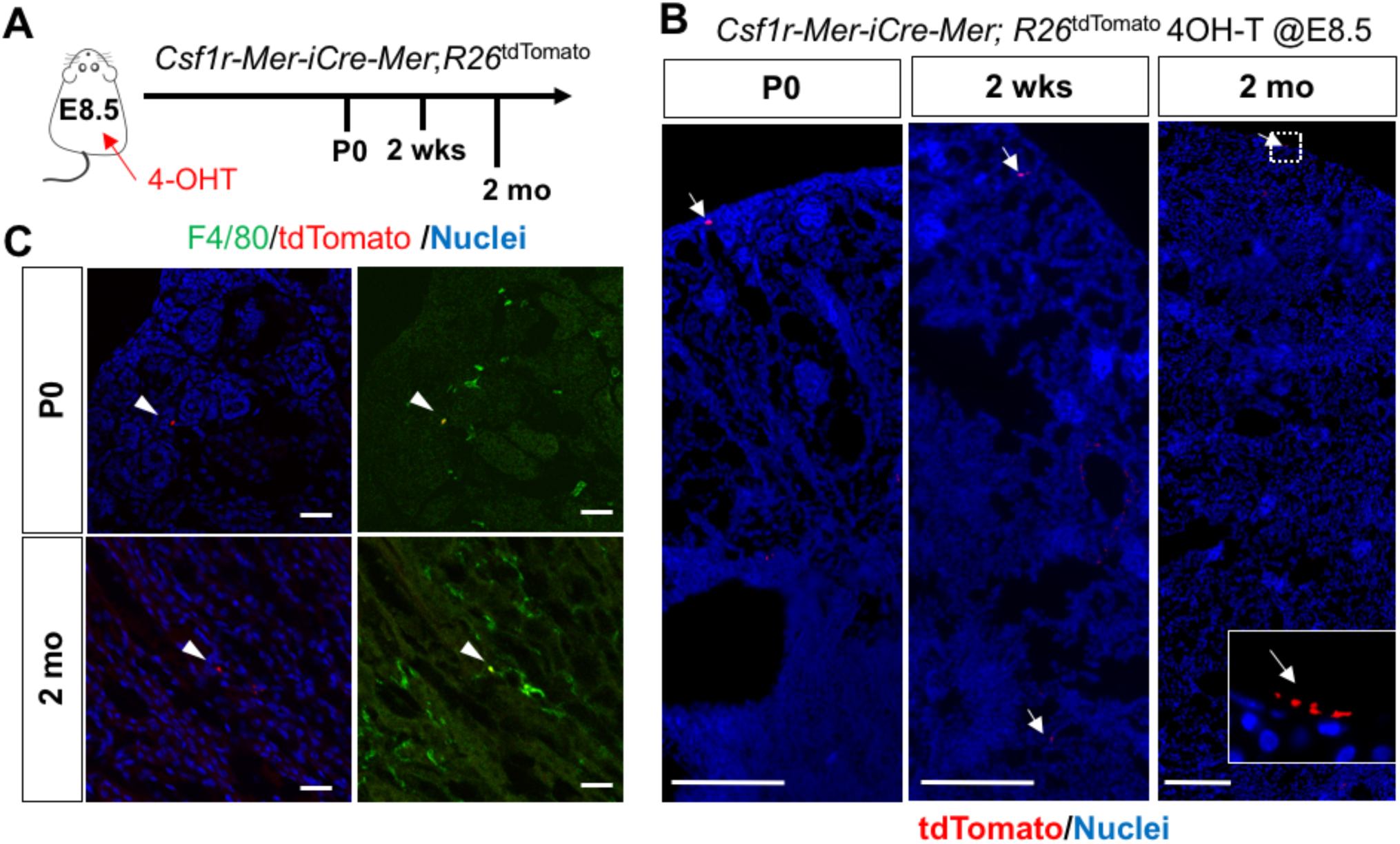
CSF1R-positive yolk-sac macrophage descendants do not expand in number in kidneys. (A) Fate-mapping strategies of CSF1R^+^ yolk-sac macrophages. 4-hydroxytamoxifen (4-OHT) was injected once into pregnant dams at 8.5 dpc and offspring analyzed at the indicated times (n=4-6). Yolk-sac macrophages and their progeny are irreversibly tagged with tdTomato. (B) Distribution of CSF1R-lineage cells in postnatal kidneys (n=4). Arrows: CSF1R-lineage cells. Inset: higher magnification of dotted box. (C) Confocal images of F4/80 staining with CSF1R-lineage tracing (n=3). Arrowheads: F4/80^+^ CX3CR1-lineage cells. Scale bars: 200 μm in B; and in 20 μm in C.

Our findings raise the question of the underlying mechanisms responsible for the chronological shift of macrophage composition. One possibility is recruitment of yolk-sac-derived macrophages from extra-renal reservoirs through the circulation. To test this hypothesis, we generated a parabiotic union between young *Cx3cr1^GFP/+^* and *Cx3cr1Cre^ER^*; *Rosa26-tdTomato* mice that had been exposed to 4-OHT *in utero* at E9.5 (Fig. 3A). Effective blood sharing between the pair was confirmed by detecting *Cx3cr1*-promoter-driven GFP expression in bone marrow cells derived from both mice (data not shown). When analyzed at 5 weeks after parabiosis, we found a few tdTomato^+^ cells in the extravascular, interstitial area of the cortex and medulla of the *Cx3Cr1*^GFP^ parabiont kidneys (Fig. 3, B and C). These results show that circulating yolk-sac macrophage descendants can slowly contribute to the adult renal macrophage pool. Currently, the origin of the circulating CX3CR1-lineage cells is not known but we speculate that one site is the spleen, which we recently identified as a reservoir of CX3CR1^+^yolk-sac macrophages (Y.Y. *in revision*).

**Figure 3.**
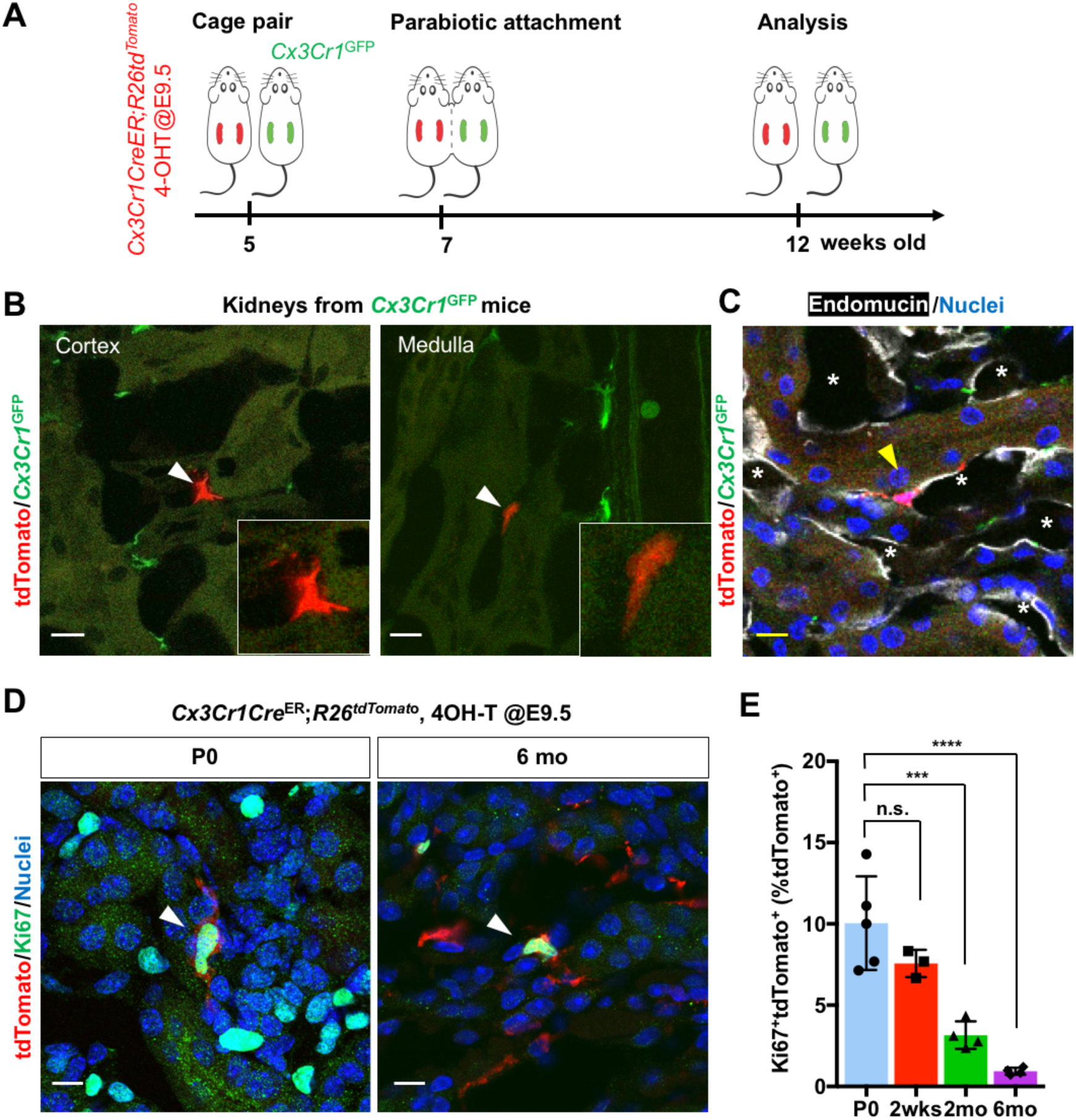
CX3CR1-positive yolk-sac macrophage descendants are recruited into adult kidneys from the circulation. (A) Schematic of parabiotic experiments. (B and C) Localization of tdTomato^+^ CX3CR1-lineage cells in *Cx3cr1*^GFP^ parabiont. Note that tdTomato^+^ cells were detected in extravascular interstitium (n = 4 per group). Endomucin; an endothelial cell marker. *, lumen of capillaries. Arrowheads, CX3CR1-lineage cells from circulation. (D and E) CX3CR1^+^ yolk-sac macrophages proliferate in neonatal and aged kidneys. *Cx3Cr1Cre^ER^; Rosa26-tdTomato* mice were treated with 4-hydroxytamoxifen (4-OHT) at E9.5 (n = 3-5). Arrowheads: Ki67^+^ CX3CR1-lineage cells. Percentage of Ki67^+^tdTomato^+^ cells to total tdTomato^+^ cells are shown in E. Data are represented as means ± S.D. ***, *P*<0.001; ****, *P*<0.0001; n.s., not significant. Scale bars: 20 μm.

We also tested whether the proliferation of yolk-sac-derived renal macrophages can potentially contribute to their population dynamics. *Cx3cr1* lineage-labeled cells that express the proliferation marker, Ki67, are present in kidneys from birth (Fig. 3, D and E). Although the percentage declines with age, Ki67^+^ cells are still present at 6 months, indicating that yolk-sac-derived macrophages retain the potential to expand in numbers through proliferation.

In conclusion, we have shown here that the proportion of yolk-sac-derived, CX3CR1-positive, macrophages increases significantly in the kidney with age, with recruitment from the circulation and proliferation being two possible mechanisms. Our findings provide a foundation for future studies to investigate the functional heterogeneity of ontogenically distinct renal macrophages in younger versus aged kidneys. These future studies may provide novel insight into age-related susceptibility of the kidney to acute and chronic diseases.

## Materials and Methods

### Study approval

All experiments were performed according to IACUC-approved protocols.

### Animals

The mouse lines were from the Jackson Laboratory (Stock No: 019098; 020940; 007914; and 005582). 75μg/g body weight of 4-hydroxytamoxifen (4-OHT; Sigma Aldrich, St. Louis, MO) dissolved in corn oil (Sigma Aldrich) was intraperitoneally administered into pregnant dams with 37.5μg/g body weight progesterone (Sigma Aldrich) to avoid fetal abortions. Mice without 4-OHT treatment were used for the specificity of tdTomato signals (Figure 1 supplement figure 1). Animals were allocated to the experimental group randomly into experimental groups and analyses. To determine experimental group sizes, data from our previous study (Y.Y in revision) were used to estimate the required numbers.

### Antibodies and Sample Processing

Primary antibodies: F4/80 (Bio-Rad, Hecules, CA; clone Cl:A3-1), CD64 (Bio-Rad; clone AT152-9), Endomucin (Abcam; Cambridge, UK; clone V.7C7.1), Ki67 (eBioscience, San Diego, CA; clone SolA15 and Thermo, Waltham, MA; clone SP6), GFP (MBL, Nagoya, Japan; cat. #598), and dsRed (Rockland, Limerick, PA; cat. #600-401-379). Fluorescent-labeled secondary antibodies were used appropriately. 7-μm cryosections were stained using standard protocols. Images were captured using Axio imager and 780 confocal microscopes (Zeiss, Oberkochen, Germany). More than 3 randomly selected areas from 3 to 5 kidneys were imaged and quantified using Image-J.

### Statistics and Reproducibility

Results are expressed as means ± SD. One-way ANOVA followed by Dunnett’s correction was used. A P value less than 0.05 was considered statistically significant.

## Acknowledgments

This study was supported by grants from the National Institute on Aging R01 AG049745 to B.A., and NIH-AI088100 to M.L.S., and the American Society of Nephrology Career Developmental Grant and Duke Nephrology Start-up Fund to T. S. We thank Drs. Brigid Hogan, Benjamin Thomson, and Myles Wolf for comments and helpful suggestions on the manuscript.

## Author contributions

S.I., Y.Y., and T.S designed research; S.I., Y.Y., S.S., A.W., J.R.P. performed research; Y. K., J.R.P., S.C., M.L.S., and B.A contributed new reagents/analytic tools; S.I., Y.Y, and T.S. analyzed data; and S.I., Y.Y, J.R.P, M.L.S., and T.S. wrote the paper

## References

1. Mass E. Delineating the origins, developmental programs and homeostatic functions of tissue-resident macrophages. Int Immunol, 2018;30(11):493–501. doi: 10.1093/intimm/dxy044.

2. Ginhoux F, Greter M, Leboeuf M, Nandi S, See P, Gokhan S, Mehler MF, Conway SJ, Ng LG, Stanley ER, Samokhvalov IM, & Merad M. Fate mapping analysis reveals that adult microglia derive from primitive macrophages. Science, 2010;330(6005):841–845. doi: 10.1126/science.1194637.

3. Schulz C, Gomez Perdiguero E, Chorro L, Szabo-Rogers H, Cagnard N, Kierdorf K, Prinz M, Wu B, Jacobsen SE, Pollard JW, Frampton J, Liu KJ, & Geissmann F. A lineage of myeloid cells independent of Myb and hematopoietic stem cells. Science, 2012;336(6077):86–90. doi: 10.1126/science.1219179.

4. Yona S, Kim KW, Wolf Y, Mildner A, Varol D, Breker M, Strauss-Ayali D, Viukov S, Guilliams M, Misharin A, Hume DA, Perlman H, Malissen B, Zelzer E, & Jung S. Fate mapping reveals origins and dynamics of monocytes and tissue macrophages under homeostasis. Immunity, 2013;38(1):79–91. doi: 10.1016/j.immuni.2012.12.001.

5. Gomez Perdiguero E, Klapproth K, Schulz C, Busch K, Azzoni E, Crozet L, Garner H, Trouillet C, de Bruijn MF, Geissmann F, & Rodewald HR. Tissue-resident macrophages originate from yolk-sac-derived erythro-myeloid progenitors. Nature, 2015;518(7540):547–551. doi: 10.1038/nature13989.

6. Mass E, Ballesteros I, Farlik M, Halbritter F, Gunther P, Crozet L, Jacome-Galarza CE, Handler K, Klughammer J, Kobayashi Y, Gomez-Perdiguero E, Schultze JL, Beyer M, Bock C, & Geissmann F. Specification of tissue-resident macrophages during organogenesis. Science, 2016;353(6304):aaf4238–aaf4238. doi: 10.1126/science.aaf4238.

7. Hashimoto D, Chow A, Noizat C, Teo P, Beasley Mary B, Leboeuf M, Becker Christian D, See P, Price J, Lucas D, Greter M, Mortha A, Boyer Scott W, Forsberg EC, Tanaka M, van Rooijen N, García-Sastre A, Stanley ER, Ginhoux F, Frenette Paul S, & Merad M. Tissue-Resident Macrophages Self-Maintain Locally throughout Adult Life with Minimal Contribution from Circulating Monocytes. Immunity, 2013;38(4):792–804. doi: 10.1016/j.immuni.2013.04.004.

8. Epelman S, Lavine KJ, Beaudin AE, Sojka DK, Carrero JA, Calderon B, Brija T, Gautier EL, Ivanov S, Satpathy AT, Schilling JD, Schwendener R, Sergin I, Razani B, Forsberg EC, Yokoyama WM, Unanue ER, Colonna M, Randolph GJ, & Mann DL. Embryonic and adult-derived resident cardiac macrophages are maintained through distinct mechanisms at steady state and during inflammation. Immunity, 2014;40(1):91–104. doi: 10.1016/j.immuni.2013.11.019.

9. Stremmel C, Schuchert R, Wagner F, Thaler R, Weinberger T, Pick R, Mass E, Ishikawa-Ankerhold HC, Margraf A, Hutter S, Vagnozzi R, Klapproth S, Frampton J, Yona S, Scheiermann C, Molkentin JD, Jeschke U, Moser M, Sperandio M, Massberg S, Geissmann F, & Schulz C. Yolk sac macrophage progenitors traffic to the embryo during defined stages of development. Nat Commun, 2018;9(1). doi: 10.1038/s41467-017-02492-2.

10. Hoeffel G, Chen J, Lavin Y, Low D, Almeida Francisca F, See P, Beaudin Anna E, Lum J, Low I, Forsberg EC, Poidinger M, Zolezzi F, Larbi A, Ng Lai G, Chan Jerry KY, Greter M, Becher B, Samokhvalov Igor M, Merad M, & Ginhoux F. C-Myb+ Erythro-Myeloid Progenitor-Derived Fetal Monocytes Give Rise to Adult Tissue-Resident Macrophages. Immunity, 2015;42(4):665–678. doi: 10.1016/j.immuni.2015.03.011.

11. Sawai CM, Babovic S, Upadhaya S, Knapp D, Lavin Y, Lau CM, Goloborodko A, Feng J, Fujisaki J, Ding L, Mirny LA, Merad M, Eaves CJ, & Reizis B. Hematopoietic Stem Cells Are the Major Source of Multilineage Hematopoiesis in Adult Animals. Immunity, 2016;45(3):597–609. doi: 10.1016/j.immuni.2016.08.007.

12. Soucie EL, Weng Z, Geirsdottir L, Molawi K, Maurizio J, Fenouil R, Mossadegh-Keller N, Gimenez G, VanHille L, Beniazza M, Favret J, Berruyer C, Perrin P, Hacohen N, Andrau JC, Ferrier P, Dubreuil P, Sidow A, & Sieweke MH. Lineage-specific enhancers activate self-renewal genes in macrophages and embryonic stem cells. Science, 2016;351(6274):aad5510. doi: 10.1126/science.aad5510.

13. Bain CC, Bravo-Blas A, Scott CL, Perdiguero EG, Geissmann F, Henri S, Malissen B, Osborne LC, Artis D, & Mowat AM. Constant replenishment from circulating monocytes maintains the macrophage pool in the intestine of adult mice. Nat Immunol, 2014;15(10):929–937. doi: 10.1038/ni.2967.

14. Zhu Y, Herndon JM, Sojka DK, Kim KW, Knolhoff BL, Zuo C, Cullinan DR, Luo J, Bearden AR, Lavine KJ, Yokoyama WM, Hawkins WG, Fields RC, Randolph GJ, & DeNardo DG. Tissue-Resident Macrophages in Pancreatic Ductal Adenocarcinoma Originate from Embryonic Hematopoiesis and Promote Tumor Progression. Immunity, 2017;47(2):323–338 e326. doi: 10.1016/j.immuni.2017.07.014.

15. Stamatiades EG, Tremblay ME, Bohm M, Crozet L, Bisht K, Kao D, Coelho C, Fan X, Yewdell WT, Davidson A, Heeger PS, Diebold S, Nimmerjahn F, & Geissmann F. Immune Monitoring of Trans-endothelial Transport by Kidney-Resident Macrophages. Cell, 2016;166(4):991–1003. doi: 10.1016/j.cell.2016.06.058.

16. Berry MR, Mathews RJ, Ferdinand JR, Jing C, Loudon KW, Wlodek E, Dennison TW, Kuper C, Neuhofer W, & Clatworthy MR. Renal Sodium Gradient Orchestrates a Dynamic Antibacterial Defense Zone. Cell, 2017;170(5):860–874 e819. doi: 10.1016/j.cell.2017.07.022.

17. Viehmann SF, Bohner AMC, Kurts C, & Brahler S. The multifaceted role of the renal mononuclear phagocyte system. Cell Immunol, 2018;330:97–104. doi: 10.1016/j.cellimm.2018.04.009.

18. Munro DAD, Wineberg Y, Tarnick J, Vink CS, Li Z, Pridans C, Dzierzak E, Kalisky T, Hohenstein P, & Davies JA. Macrophages restrict the nephrogenic field and promote endothelial connections during kidney development. Elife, 2019;8. doi: 10.7554/eLife.43271.

19. Lever JM, Hull TD, Boddu R, Pepin ME, Black LM, Adedoyin OO, Yang Z, Traylor AM, Jiang Y, Li Z, Peabody JE, Eckenrode HE, Crossman DK, Crowley MR, Bolisetty S, Zimmerman KA, Wende AR, Mrug M, Yoder BK, Agarwal A, & George JF. Resident macrophages reprogram toward a developmental state after acute kidney injury. JCI Insight, 2019;4(2). doi: 10.1172/jci.insight.125503.

20. Chen T, Cao Q, Wang Y, & Harris DCH. M2 macrophages in kidney disease: biology, therapies, and perspectives. Kidney Int, 2019;95(4):760–773. doi: 10.1016/j.kint.2018.10.041.

21. Chawla LS, Eggers PW, Star RA, & Kimmel PL. Acute Kidney Injury and Chronic Kidney Disease as Interconnected Syndromes. N Engl J Med, 2014;371(1):58–66. doi: 10.1056/NEJMra1214243.

22. Sato Y & Yanagita M. Immunology of the ageing kidney. Nat Rev Nephrol, 2019. doi: 10.1038/s41581-019-0185-9.

23. Sato Y, Mii A, Hamazaki Y, Fujita H, Nakata H, Masuda K, Nishiyama S, Shibuya S, Haga H, Ogawa O, Shimizu A, Narumiya S, Kaisho T, Arita M, Yanagisawa M, Miyasaka M, Sharma K, Minato N, Kawamoto H, & Yanagita M. Heterogeneous fibroblasts underlie age-dependent tertiary lymphoid tissues in the kidney. JCI Insight, 2016;1(11):e87680. doi: 10.1172/jci.insight.87680.

24. Molawi K, Wolf Y, Kandalla PK, Favret J, Hagemeyer N, Frenzel K, Pinto AR, Klapproth K, Henri S, Malissen B, Rodewald HR, Rosenthal NA, Bajenoff M, Prinz M, Jung S, & Sieweke MH. Progressive replacement of embryo-derived cardiac macrophages with age. J Exp Med, 2014;211(11):2151–2158. doi: 10.1084/jem.20140639.

25. Bain CC, Hawley CA, Garner H, Scott CL, Schridde A, Steers NJ, Mack M, Joshi A, Guilliams M, Mowat AM, Geissmann F, & Jenkins SJ. Long-lived self-renewing bone marrow-derived macrophages displace embryo-derived cells to inhabit adult serous cavities. Nat Commun, 2016;7:ncomms11852. doi: 10.1038/ncomms11852.

26. Dick SA, Macklin JA, Nejat S, Momen A, Clemente-Casares X, Althagafi MG, Chen J, Kantores C, Hosseinzadeh S, Aronoff L, Wong A, Zaman R, Barbu I, Besla R, Lavine KJ, Razani B, Ginhoux F, Husain M, Cybulsky MI, Robbins CS, & Epelman S. Self-renewing resident cardiac macrophages limit adverse remodeling following myocardial infarction. Nat Immunol, 2019;20(1):29–39. doi: 10.1038/s41590-018-0272-2.

27. Brahler S, Zinselmeyer BH, Raju S, Nitschke M, Suleiman H, Saunders BT, Johnson MW, Bohner AMC, Viehmann SF, Theisen DJ, Kretzer NM, Briseno CG, Zaitsev K, Ornatsky O, Chang Q, Carrero JA, Kopp JB, Artyomov MN, Kurts C, Murphy KM, Miner JH, & Shaw AS. Opposing Roles of Dendritic Cell Subsets in Experimental GN. J Am Soc Nephrol, 2018;29(1):138–154. doi: 10.1681/ASN.2017030270.

