## Supplement Figure 1 for "Yolk-Sac-Derived Macrophages Progressively Expand in the Mouse Kidney with Age"

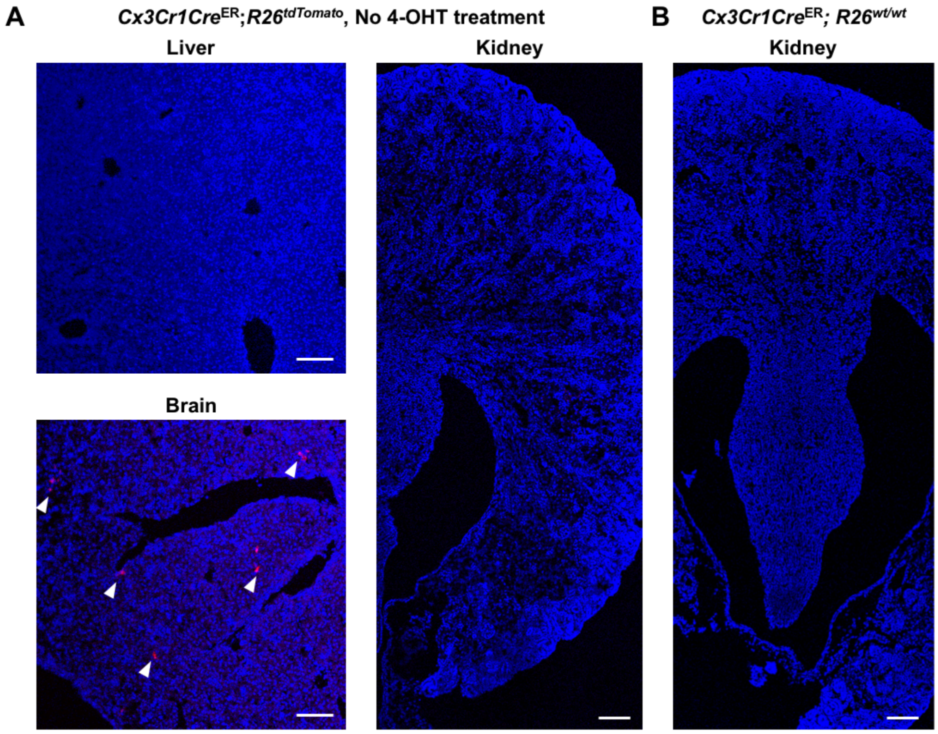


**Figure 1 supplement figure 1.** There is no basal Cre activity in kidneys without 4-hydroxytamoxifen (4-OHT) treatment. *Cx3cr1Cre*$^{ER}$; *R26*$^{\mathrm{tdTomato/wt}}$ mice were used to determine tdTomato reporter activity in the absence of 4-OHT treatment. (A) While we observed a few tdTomato-expressing cells in the brain, there were no tdTomato-positive cells in the liver and kidney at P3 without 4-OHT treatment. Scale bars: 100 µm. (B) *Cx3cr1Cre*$^{ER}$; *R26*$^{\mathrm{wt/wt}}$ mice from the same litter was used as a negative control. Arrowheads: tdTomato-expressing cells.
